# Combining transformer and 3DCNN models to achieve co-design of structures and sequences of antibodies in a diffusional manner

**DOI:** 10.1101/2024.04.25.587828

**Authors:** Yue Hu, Feng Tao, WenJun Lan, Jing Zhang

## Abstract

Antibody drugs are among the fastest growing therapeutic modalities in modern drug research and development. Due to the huge search space of antibody sequences, the traditional experimental screening method cannot fully meet the needs of antibody discover. More and more rational design methods have been proposed to improve the success rate of antibody drugs. In recent years, artificial intelligence methods have increasingly become an important means of rational design. We have proposed an algorithm for antibody design, called AlphaPanda (AlphaFold2 inspired Protein-specific antibody design in a diffusional manner). The algorithm mainly combines the transformer model, the 3DCNN model and the diffusion generative model, use the transformer model to capture the global information and uses the 3DCNN model to capture the local structural characteristics of the antibody-antigen complexes, and then uses the diffusion model to generate sequences and structures of antibodies. The 3DCNN model can capture pairwise interactions in antibody-antigen complex, as well as non-pairwise interactions in antibody-antigen complex, and it requires less training sample data, while avoiding the defects of the generation progress by the autoregressive model and by the self-consistent iterative model. Diffusion generative model can generate sequence and structure effectively and with high quality. By combining 3DCNN method and diffusion model method, we have achieved the integration of 3DCNN model to the protein design with flexible main chains. By utilizing the advantages of these aspects, a good performance has been achieved by the AlphaPanda algorithm. The algorithm we propose can not only be applied to antibody design, but also be more widely applied to various fields of other protein design. The source code can be get from github (https://github.com/YueHuLab/AlphaPanda).

## 1. Introduction

Biological drugs represented by monoclonal antibodies have become a hot spot in drug research and development in recent year^1-2^. From 1986, the first monoclonal antibody was approved by the US Food and Drug Administration (FDA), and by 2023, the first sale of the monoclonal antibody drug (Keytruda) exceeded 25 billion US dollars. With the development of monoclonal antibodies, a series of technologies have emerged, including humanized antibody modification technology^3^, phage display^4-5^, humanized antibody mouse^6^, single B cell antibody technology^7^, fully synthetic antibody gene bank technology^8^ and so on.

Although these are all high-throughput experimental methods, their throughput is still insufficient relative to the sequence space of antibodies. It is necessary to further reduce the screening space by combining rational design to improve the success rate of antibody drug design^9-10^.

There are many protein and antibody design programs^11-12^, such as diffAb^13^, PROTSEED^14^, Rosetta^15-17^, RFdiffusion^18-19^, SCUBA^20^, Abacus-R^21^, Fold-X^22^, EGAD^23-24^, 3DCNN^25-26^ etc. All of these have the potential to design antibodies targeting specific epitopes. At present, the deep learning protein design method can overcome the shortcomings of the traditional energy function method (the energy surface is rugged and trapped in local minima), and can process high-dimensional spatial data. It uses many advanced methods^27^, such as discriminative models and generative models^28^, convolutional neural network models, recurrent neural network models^29^, adversarial network models^28^, variational auto-encoders^30^, transformer model^31-32^, graph neural network model^33-34^ and so on.

At present, there are still some important aspects to be considered for designing antibodies based on deep learning: 1, How to capture both global and local information simultaneously? At the same time, effective training can be conducted for limited antibody antigen structure data. 2,How can we capture both the pair contacts and the high-dimensional unpair contacts for antigen antibody complexes? 3, How to efficiently generate the structure and sequence of antibodies? In the generation mode of autoregression, the upstream errors tend to accumulate in the downstream errors. The generation method of self-consistent iteration does not have an effective function to guide the direction of generation. 4, How to use the 3DCNN method to design proteins for flexible main chains?

In particular, the methods to explore the above aspects, but there are still some shortcomings. The diffab method uses diffusion model to co-generate the sequence and structure of antibody CDR region. The PROTSEED method adds geometric constraints and provides the interaction of information processing, but does not adopt a high-quality generation model. The above two methods do not focus on local information or consider the unpair contacts. The 3DCNN method did not consider global information and did not adopt an efficient generation method. RFdiffusion divides antibody design into two stages, first designing the structure of the antibody, and then designing the sequence of the antibody.

Inspired by AlphaFold2^35-36^ and using diffusion model, we construct AlphaPanda (AlphaFold2 inspired Protein-specific antibody design in a diffusional manner) algorithm. In this model, we will combine the Transformer model with the 3DCNN model. And a diffusion model^37^ is used to generate sequences and structures. In this way, we consider both the overall information and the global information, taking into account both the pair contacts and the unpair contact, while avoiding the defects of the generation progress by the autoregressive model and by the self-consistent iterative model. By combining 3DCNN method and diffusion model method, we have achieved the integration of 3DCNN model to the protein design with flexible main chains.

Transformers and CNNs (Convolutional Neural Networks) are powerful models used in various applications. Transformers can capture long-range dependencies, while CNNs are excellent at learning local patterns. Combining them can lead to a more robust feature representation. The combination can leverage the strengths of both models, potentially outperforming each used individually in certain tasks^38^. Please note that the combined model can be more complex and require more computational resources. Transformers often need large amounts of data to perform well, which might not be necessary for CNNs. This is also why introducing CNNs to learn antibody structures may be easier to converge.

Combining Transformer and CNN models can elevate the properties of a model by leveraging the strengths of both architectures in many fields. For example, the ConViT model in vision tasks^39^. A model that combines Transformers and CNNs to overcome limitations of each approach on its own. It performs well in low data regimes and achieves similar performance in large data settings. And in image classification area, the CofaNet model^40^, a classification network that combines CNNs and transformer-based fused attention. It introduces patch sequence dimension attention to capture the relationship among subsequences and incorporates it into self-attention to construct a new attention feature extraction layer. These models provide detailed insights into how the combined use of Transformers and CNNs can enhance model performance by integrating local and global feature extraction capabilities. Herein, we combined those two models in antibody design area.

We implemented our AlphaPanda based on the diffab program and the 3DCNN program. Therefore, AlphaPanda algorithm has the common advantages of other advanced protein or antibody design software. It uses equivariant neural network to deal with coordinates in three-dimensional space, which ensures equivariant. It explicitly considers the structure of antigen and realizes the simultaneous diffusion generation of sequence and structure of CDR region. And AlphaPanda is an end-to-end generation model.

By testing our method, we have achieved good results in various indicators. This method is not limited to antibodies design (or nanobodies design^41-47^), and can be widely used in other fields of protein design (fixed backbone design, antibody optimization, etc.).

## 2. Method

### 2.1 The overall of AlphaPanda

Inspired by AlphaFold2 and other protein design methods combined with diffusion generation model, we propose AlphaPanda algorithm. We have written AlphaPanda based diffab and 3DCNN, which were written by python and pytorch. Because diffab program is mainly for antibody design, we added the 3DCNN program as a module to diffab. The software is open source and licensed by Apache 2.0. Everyone can get the source code from github (***https://github.com/YueHuLab/AlphaPanda***).

The diffab is a cutting-edge tool for antigen-specific antibody design and optimization, utilizing diffusion-based generative models for protein structures. It was presented at NeurIPS 2022 and is notable for being one of the first deep learning-based methods that generate antibodies targeting specific antigen structures. The diffab model operates by jointly modeling the sequences and structures of complementarity-determining regions (CDRs) of antibodies. These CDRs are crucial as they determine the binding affinity between antibodies and antigens, such as viruses and bacteria. By leveraging diffusion probabilistic models and equivariant neural networks, diffab can perform several tasks: Sequence-structure co-design, Sequence design, Structure prediction and Antibody optimization. The diffab is designed to be a versatile ‘Swiss Army Knife’ for antibody design, capable of addressing various challenges in the field. The model has been evaluated extensively, showing promising results in terms of biophysical energy functions and other protein design metrics. The software using transformer model to get the information of the antigen-antibody complexes. The 3DCNN protein design software is a computational tool developed by Anand Namrata, which utilizes a 3D Convolutional Neural Network (3DCNN) for analyzing protein structures and protein design. This tool represents an innovative approach to protein structure analysis, leveraging deep learning to gain insights into the complex world of proteins.

We combined that two softwares, utilized the transformer and 3DCNN model to embedding the information of the antigen-antibody complexes. Combining diffab and 3DCNN could potentially offer several advantages:3DCNN’s ability to analyze protein structures could complement diffab’s design capabilities, providing a more comprehensive understanding of protein interactions; The structural insights from 3DCNN could inform diffab’s generative models, potentially leading to more accurate and efficient antibody designs; The combination could lead to a broader range of applications, from antibody design to broader protein engineering tasks. Overall, the integration of these two software tools could lead to a powerful platform for protein engineering, expanding the possibilities for research and development in the field.

The following is a brief introduction to the main ideas and algorithms of AlphaPanda. Proteins are made up of amino acid residues. An amino acid residue can be represented by its sequence type*s*_*i*_ ∈ {ACDEFGHIKLMNPQRSTVWY}, its atomic coordinates of alpha-carbon C_*α*_***x***_*i*_ ∈ ℝ^3^ and its frame orientation ***O***_*i*_ ∈ SO(^3^) respectively. The latter two items are structural information. So we represent both sequence and structural information simultaneously in this way.

In (nano) antibody design, it is required to target antigen epitopes, so the general position of antigen-antibody complex binding is determined. The main problem is to generate the sequence and structure information of CDR region of antibody by deep learning method given the remaining antigen-antibody complex structure and sequence information. We represent the given information of protein and the information sampled in the generation iteration process as context features. The environmental information is embedded using the transformer model and the 3DCNN model. The transformer model is expressed by single base information (including residue type, secondary structure information, three-dimensional coordinates of heavy atoms, etc.) and base pair information (pairwise interaction information, pairwise spatial distance information, etc.). The 3DCNN model is mainly considering information about the position of surrounding atoms. 3DCNN model does not consider protein chains, but only considers the spatial position of atoms, which may make interaction design, especially for long-chain and multi-chain proteins, simpler.

Dependent on environmental information, the structure and sequence of CDR regions 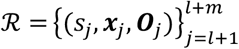 are generated by generation diffusion iteration. Together with known ones, the structural information of the remaining protein antigen and antibody complexes 𝒞 = {(*s*_*i*_, ***x***_*i*_, ***O***_*i*_)|*i* ∈ {1 . . . *N*}\{*l* + 1, . . ., *l* + *m*}}, constitutes the amino acid information of the entire protein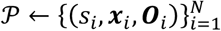.

The generated information of these amino acids of proteins 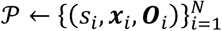 is transmitted through a IPA like module of Alphafold2 into the Transformer model. Meanwhile, the generated information 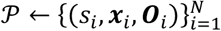 is also transmitted to the 3DCNN model by updating the protein structure and sequence.

## The overall flow is shown in Algorithm 1

### Algorithm 1

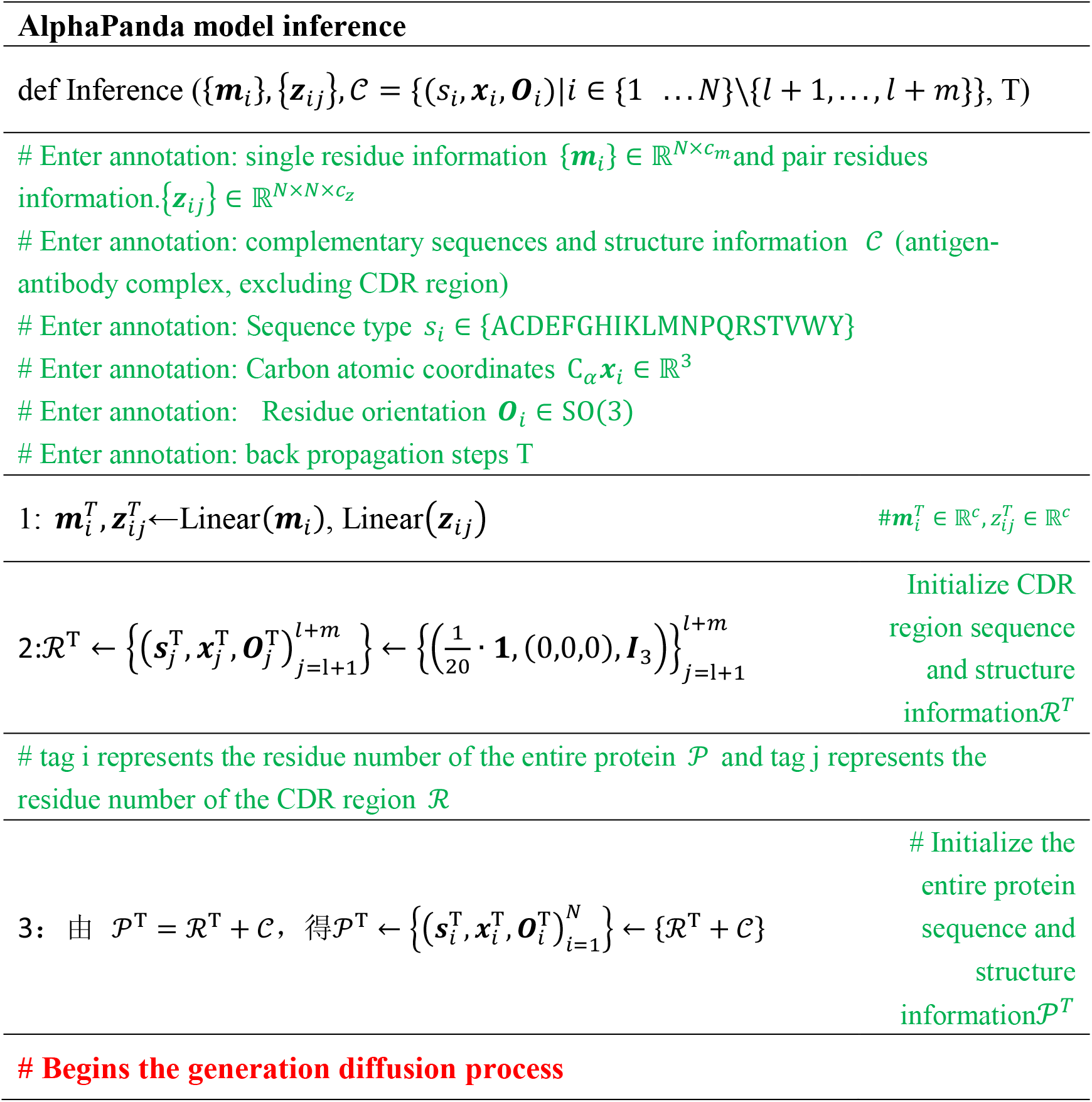
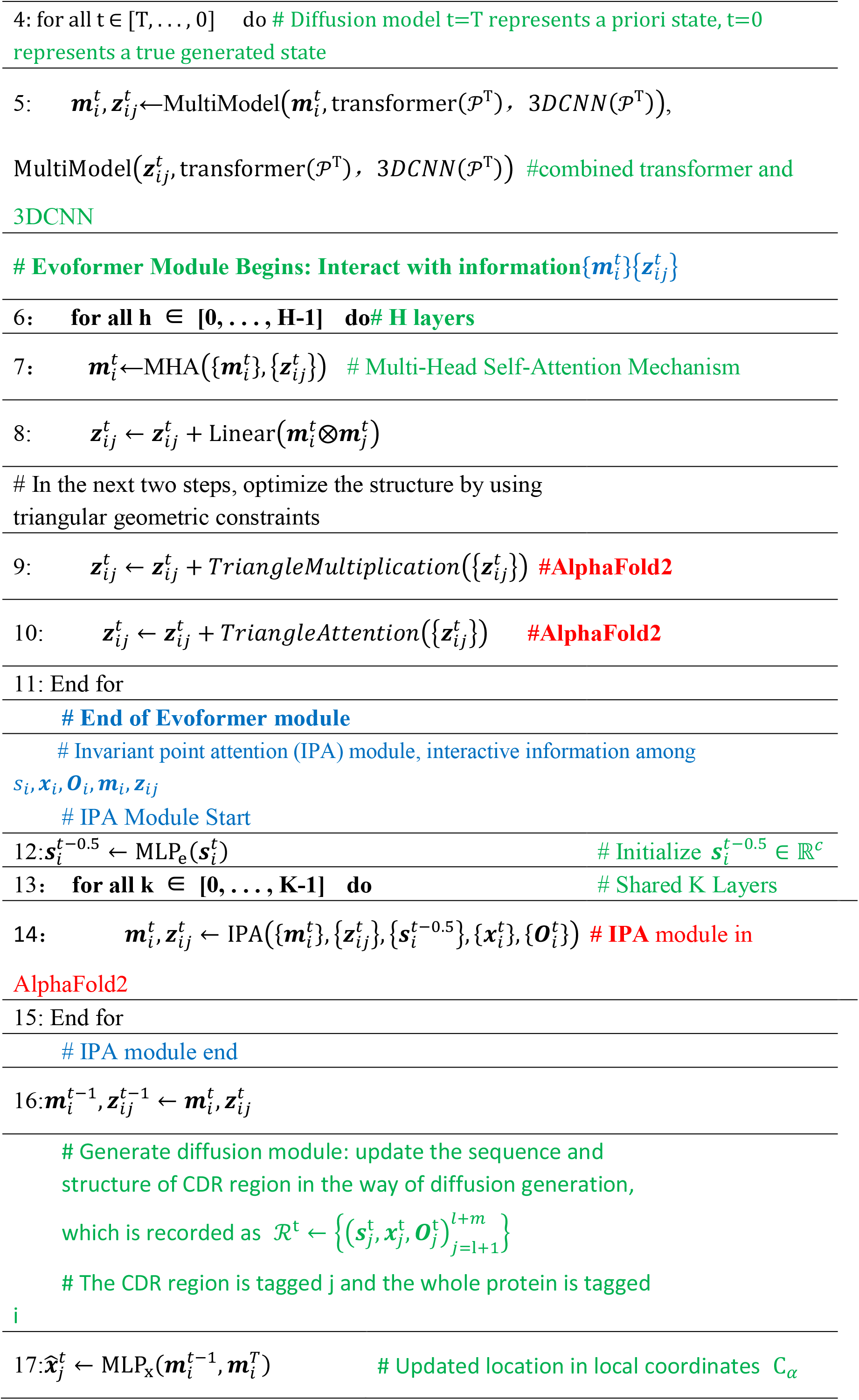
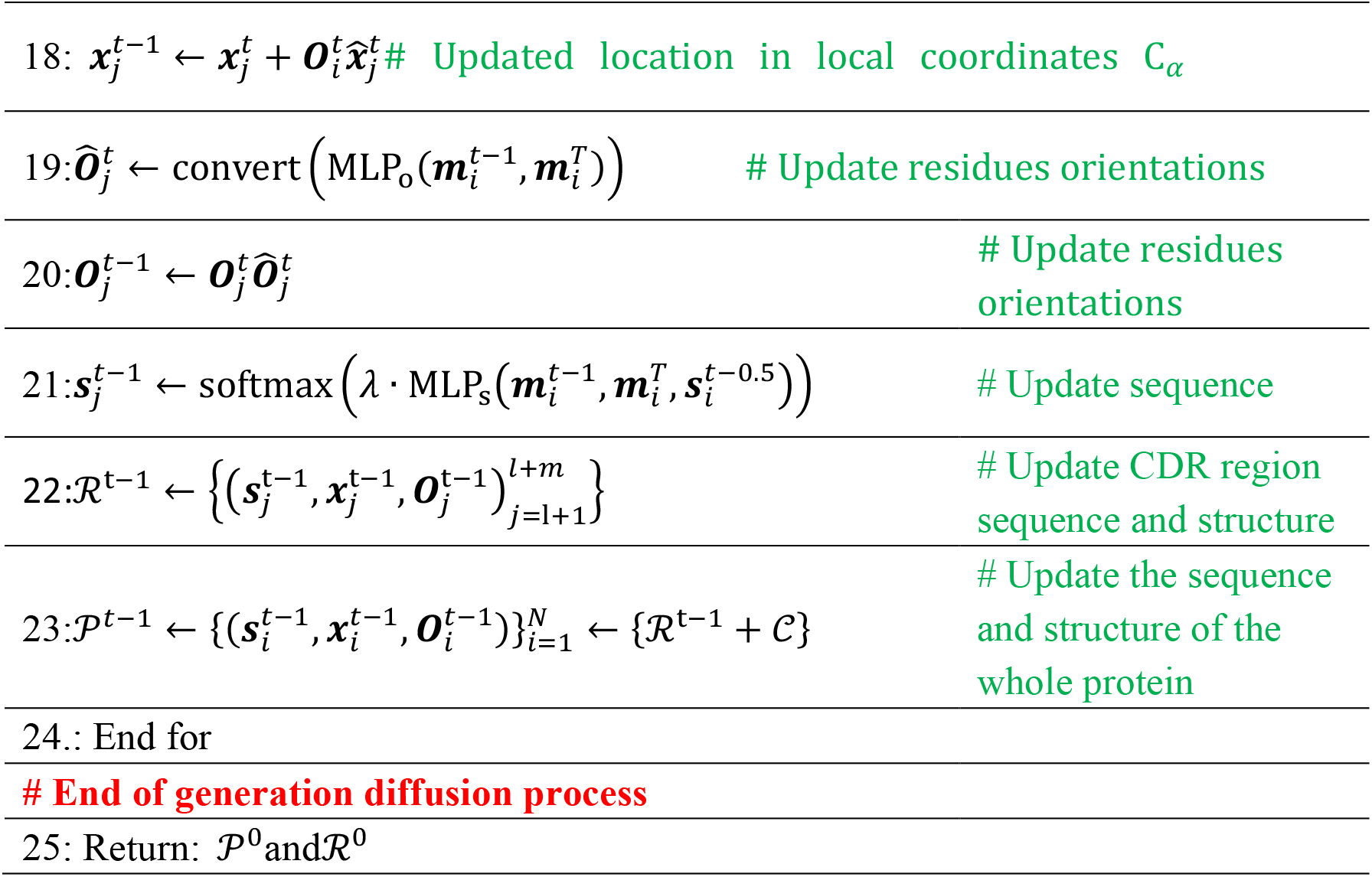

### 2.2 Training details

Diffusion probability model is a common generation model, which includes two Markov processes: the positive process gradually adds noise, which makes the data distribution (real distribution) gradually change into a prior distribution (generally Gaussian distribution); In the reverse generation process, the prior distribution is denoised to achieve the desired distribution (generating the real distribution). It regards the generation process as a step-by-step denoising process. Training the process of generation diffusion (reverse generation process) to estimate the distribution P can achieve the purpose of protein sequence-structure co-generation. This process requires the forward process to calculate the posterior distribution q (posterior). Then, different objective functions are used to estimate the difference between p and q, and the training model is optimized.

We denote the sequence structure state of the entire CDR region by representing the state of amino acid residue j at step T. The step t=0 is recorded as the true state (CDR region sequence and structure of natural protein are used in training), and t=T is recorded as a priori state distribution (generally Gaussian distribution). Forward process from t=0 to t=T; The reverse generation process ranges from t=T to t=0. The implementation of diffusion model is introduced below. I is used as the residue serial number of the whole complex, and J is used to represent the residue serial number of antibody CDR region.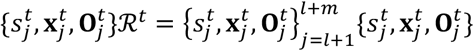

A.Diffusion Model of Amino Acid Types

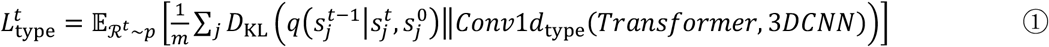

B.Diffusion model of atomic coordinatesC_*α*_***x***_*i*_

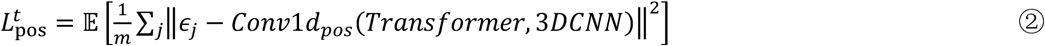

C.Frame Orientation Diffusion Process of Amino Acid***O***_*i*_ ∈ SO(^3^)

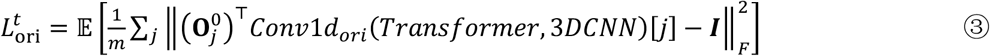

D.Training process

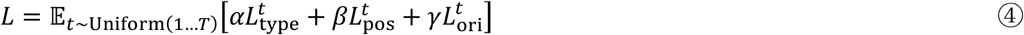

In the training process, we first select a t, then calculate the posterior probability q, estimate the posterior probability p with the corresponding MLP network, and calculate the difference loss between them using above equations. Back propagate loss to update model parameters (the whole network, including protein sequence language model, is fine-tuned) until convergence. In the training process, we first use protein-protein interaction complex structure database to train, then use antigen-antibody structure database to train, and finally use nano-antibody antigen structure database to fine-tuning.

### 2.3 Residue Orientations

The orientations of the amino acid residues in the two software are different. Here, we unified the orientations of the residues to the orientations of the residues in the diffab program.

### 2.4 Evaluated the results of the design

The root mean square deviation (RMSD), amino acid recovery (AAR), and energy (by Rosetta software) were used as criteria for comparison with other methods. We selected the antibody-antigen protein complex (PDB ID: 7xjf), specifically the crystal structure of 6MW321 1 Fab in complex with CD47, as our test case. Notably, this complex was released on May 31, 2023, and is not part of our training or test database. To evaluate our method, we designed 500 H_CDR3 sequences using the AlphaPanda software. This allowed us to assess both performance and stability. As a control, we also designed 100 H _CDR3 sequences using diffab.

## 3. Results

In our research, we explored the use of the AlphaPanda software for antibody design, which has demonstrated relatively high stability. Out of the 500 antibodies we designed, only two exhibited particularly large RMSD values. Upon closer examination, we identified the two structures that appears to be poorly designed. To further evaluate our results, we conducted a detailed comparison between these 498 structures and the 100 structures designed using diffab. Notably, the RMSD achieved by AlphaPanda was significantly (Table 1, P-value is very small and in Supplemental Table 1) about 0.5 angstrom lower than that achieved by diffab. However, in terms of energy and natural sequences consistence, there was no significant difference (Supplemental Table 1). In summary, while AlphaPanda has improved its structure generation, it remains comparable to diffab in other aspects. Details can be found in Supplemental Table 2.

**Table 1.** Group Statistics.

| **Table 1. Group Statistics** |  |  |  |  |  |
| --- | --- | --- | --- | --- | --- |
|  | class | N | Mean | Std. Deviation | Std. Error Mean |
| rmsd | 1 | 100 | 4.8684562 | 1.12474075 | .11247408 |
|  | 2 | 498 | 4.3064585 | .72066744 | .03229388 |
| seqid | 1 | 100 | 19.1949774 | 9.71249534 | .97124953 |
|  | 2 | 498 | 18.2663087 | 8.93339276 | .40031490 |
| dG_gen | 1 | 100 | 2.9722441E3 | 2.77173035E3 | 2.77173035E2 |
|  | 2 | 498 | 3.7657162E3 | 2.63051979E3 | 1.17876410E2 |
| dG_ref | 1 | 100 | 45.9770503 | .86809783 | .08680978 |
|  | 2 | 498 | 37.1638822 | .96404423 | .04319985 |

Class 1 is designed by diffab, while class 2 is designed by AlphaPanda.

## 4. Discussion

For many of the antibody design programs, they are trying to use this protein language model, or antibody language model. However, we know the protein language model or the antibody model, it actually uses this kind of coevolutionary information. But for the pathogen and the host, the antigen and the antibody, they should not be a kind of coevolutionary information. Instead, it’s a kind of negative coevolutionary, which means that the pathogen is going to escape from the direction of evolution antibody sequence. It’s possible that the protein language model can provide some of this information, but a lot of information, it still can’t provide. This is also why the latest achievement of David Baker’s design by RFdiffusion, which is obtained through experimentation, has a success rate of only 1%.

In this article, we mainly use structural information to design antigens and antibodies, which has a certain significance. However, there are also some limitations to this approach. In our future work, we may consider incorporating physical properties to capture more information. We are also exploring further attempts in this regard.

## Acknowledgements

This work was supported by Shandong Provincial Key Laboratory of Microbial Engineering (SME) and was supported by Shandong Provincial Natural Science Foundation Committee (Grant No. ZR2016HB54).

