## Supplemental Table 1 for "Combining transformer and 3DCNN models to achieve co-design of structures and sequences of antibodies in a diffusional manner"

| **Supplemental Table 1. Independent Samples Test** | | | | | | | | | | |
| --- | --- | --- | --- | --- | --- | --- | --- | --- | --- | --- |
|  |  | Levene's Test for Equality of Variances | | t-test for Equality of Means | | | | | | |
|  |  |  | |  | | | | | 95% Confidence Interval of the Difference | |
|  |  | F | Sig. | t | df | Sig. (2-tailed) | Mean Difference | Std. Error Difference | Lower | Upper |
| rmsd | Equal variances assumed | 41.459 | .000 | 6.395 | 596 | .000 | .56199774 | .08788550 | .38939481 | .73460066 |
|  | Equal variances not assumed |  |  | 4.803 | 115.839 | .000 | .56199774 | .11701843 | .33022460 | .79377087 |
| seqid | Equal variances assumed | 1.057 | .304 | .935 | 596 | .350 | .92866877 | .99362120 | -1.02275584 | 2.88009337 |
|  | Equal variances not assumed |  |  | .884 | 134.719 | .378 | .92866877 | 1.05051305 | -1.14896200 | 3.00629954 |
| dG_gen | Equal variances assumed | .418 | .518 | -2.728 | 596 | .007 | -7.93472047E2 | 2.90882793E2 | -1.36475197E3 | -2.22192128E2 |
|  | Equal variances not assumed |  |  | -2.634 | 137.156 | .009 | -7.93472047E2 | 3.01197177E2 | -1.38906272E3 | -1.97881377E2 |
| dG_ref | Equal variances assumed | 1.338 | .248 | 84.768 | 596 | .000 | 8.81316803 | .10396832 | 8.60897922 | 9.01735684 |
|  | Equal variances not assumed |  |  | 90.890 | 152.245 | .000 | 8.81316803 | .09696477 | 8.62159781 | 9.00473826 |
| ddG | Equal variances assumed | .418 | .518 | -2.758 | 596 | .006 | -8.02285216E2 | 2.90880946E2 | -1.37356151E3 | -2.31008924E2 |
|  | Equal variances not assumed |  |  | -2.664 | 137.158 | .009 | -8.02285216E2 | 3.01190012E2 | -1.39786164E3 | -2.06708795E2 |
